# Bibliometric and Geographical Analysis of Cell Death Related Literature

**DOI:** 10.1101/035204

**Authors:** Vijaykumar Yogesh Muley, Anne Hahn, Pravin Paikrao

## Abstract

Natural language processing continues to gain importance in a thriving scientific community that communicates its latest results in such a frequency that following up on the most recent developments even in a specific field cannot be managed by human readers alone. Here we summarize and compare the publishing activity of the previous years on a distinct topic across several countries, addressing not only publishing frequency and history, but also stylistic characteristics that are accessible by means of natural language processing. Though there are no profound differences in the sentence lengths or lexical diversity among different countries, writing styles approached by Part-Of-Speech tagging are similar among countries that share history or official language or those are spatially close.

## 1 Introduction

The last two decades have witnessed tremendous progress in the digitization of textual information and making them available on world wide web. At the same time, computational techniques to extract relevant information from resources scattered all over the web have been continuously developed. The scientific field that is dedicated to perform this task, among others, for example machine translation, is called natural language processing (NLP), and is emerging as important field in information retrieval to best utilize the myriads of text data available [1]. NLP can roughly be divided into sentence segmentation, tokenization, part-of-speech tagging and information retrieval. None of these tasks are trivial for general English literature, and while handling biomedical literature, pose several unique challenges.

To extract specific information from a given text, the very basic task is to demarcate sentences in a process called sentence segmentation. Tokenization is the task of splitting an input into individual words and, depending on the specific case, other units, such as punctuation marks. The tokens in a sentence then are used for Part-of-Speech tagging where the order of tokens, their immediate context and grammatical rules are taken into account and which can be analyzed for syntactical parsing of information [2, 3]; or the tokens in the document are used as bags of words where the frequencies of tokens are used for further analysis [4, 5].

Biomedical texts are characterized by increasingly complex contents, large digital supplements, and a staggering volume of publications. Thousands of research articles are available even on specialized topics and for modern scientists it is no longer possible to go through each and every document. But by using the nascent technology of NLP one can automate the information-gathering stage – leaving to human minds the more challenging activity of higher thinking and creative processes.

The PubMed database maintained at the United States National Library of Medicine (NLM) maintains millions of abstracts [6]. These abstracts are written in English by native as well as non-native English speaking authors. We aim to determine in this study whether the official language of the countries of non-native speakers has influence on their English writing.

Our results based on a corpus related to cell death and survival as topic of interest indeed support this hypothesis and off-shoot of this study, we also provide several interesting facts about cell death research for scientific community.

## 2 Methods

### Text corpus assembly

RISmed package for R statistical programming language was used to search and retrieve abstracts of review articles available in English in PubMed [6] using the following query “((cell AND growth) OR (cell AND death) OR autophagy OR apoptosis OR necrosis OR (cell AND survival) OR senescence OR aging) AND (Review[ptyp] AND hasabstract[text] AND English[lang] AND medline[sb])”

The 188,410 abstracts obtained related to cellular death and survival were preprocessed using custom Perl scripts to remove section headers which are unlikely to be part of the sentence structure. Also, some of the articles with duplicate sentences within the same abstract were removed, when they occurred directly subsequent to each other due to HTML tags.

### Mapping countries to documents

Out of those abstracts, 40,877 did not contain affiliation information; the remaining records were assigned to the countries from which the article was submitted. For that, custom Perl scripts were used to map the affiliations to specific countries; a list of which, including cities, was derived from the Maps R package and compared against the individual words of the affiliations. Different names for the same countries had to be unified (e.g. “Netherlands”, “The Netherlands”, “Netherland”). A fraction of abstracts (n=20) was inspected manually to confirm correct mapping.

### Annotating text with part-of-speech and lemma information

The TreeTagger software was used to separate out sentences and annotate words within sentences with part-of-speech tags in each abstract, indicating grammatical function [7]. TreeTagger also provides lemmas of different forms of words with the same meaning. The results are based on lemmas or original word forms throughout this study unless mentioned otherwise.

For further processing, punctuations and numbers were removed from the abstracts, and text was set into lowercase using the tm R package [8].

### Estimating lexical diversity

In order to compute the richness of vocabulary used in abstracts from each country, we randomly sampled 100 lemmatized abstracts and, calculated the lexical diversity in each by dividing the number of unique lemmas (types) with the total number of lemmatized words (tokens) in the abstract. This was repeated for hundred times and average is reported in the Table 1.

### Clustering of countries based on part-of-speech usage in writings

For each country having more than hundred abstracts, we calculated the frequency of 36 standard part-of-speech tags in each abstract, which was normalized against the number of tokens therein. The average country-wise frequency profiles were then used to perform hierarchical clustering using Euclidean distance metric and complete linkage method.

## 3 Results and Discussion

### Descriptive statistics of the corpus

Out of 188,410 abstracts retrieved by searching key words related to cell death and survival, 188,177 abstracts were analyzed. The remaining articles lacked abstracts and were not considered for further analysis. The abstracts considered were published before 2015 and date back as far as 1963. 2013 was the year with the highest number of publications, whereas in 1963 only one article was published [9]. As shown in Fig 1a, the number of yearly publications of review articles on the topic of cell death and survival has increased dramatically in the last ten years, whereas from 1963 to 1990 less than 200 reviews were published yearly. The exponential increase in publications in the last ten to twenty years may be due to the development of many experimental techniques and information technology which provides a platform for rapid communication among researchers, easy access to literature, and online submission systems adopted by most journals, which could also be a major contributory factor.

**Fig 1.**
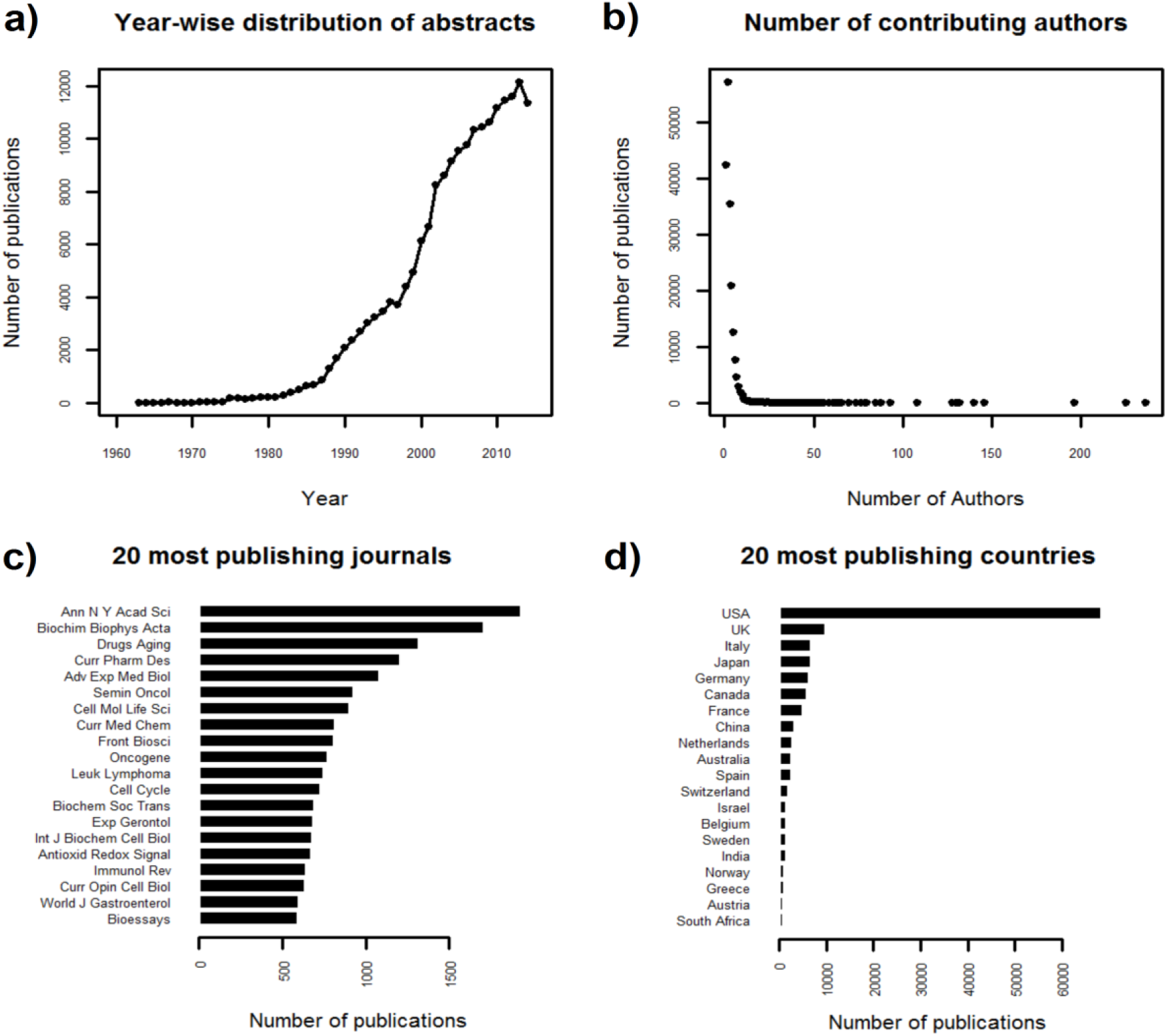
Distribution of cell death and survival related publications. A) year-wise, showing a notable increase in the past two decades. B) according to number of contributing authors. Most abstracts are published by only few authors, leaving large author consortiums the exception. C) journal-wise, in most-publishing journals. Rather than topic-specificity, a long publishing history characterizes the most publishing journals. D) country-wise, across most publishing countries. Publishing of reviews addressing cell death is dominated by the USA, in comparison, African and South American countries contributed very few abstracts and are hardly represented among the most publishing countries.

Generally, review articles are published by experts in the field and as expected we observed that most articles are published by less than five authors. However, we also observed many articles are published from more than ten up to 236 contributing authors (Fig. 1b). Manual inspection of some papers with a multitude of authors suggests that those are published by a subject specific consortium and generally are sets of guidelines, protocols, or nation-wide studies [10, 11].

The list of most publishing journals reflects the broad spectrum of subjects addressed by the selected reviews, in contrary to the expectation of journals focusing to cell death being predominant (Fig. 1c). The analysis of the top 20 most publishing journals out of 5496 suggests that rather than subject-specificity, a long publishing history of a journal is associated also to a high number of articles on the selected topics, for example Oncology or Annals of the New York Academy of Sciences (Fig. 1c). The leading journals very likely provide the reader with a comprehensive view on the topic and should be referred to for literature review.

There is no straight forward way to obtain the names of countries and cities from where articles were published. Articles were assigned to 143 countries by matching affiliation information first to countries and then to cities. By this strategy ambiguous place names could be mapped. Therefore, we analyzed 20 randomly selected abstracts and their affiliation information and found correct mapping of the respective countries and cities by our strategy. The distribution of articles across the most publishing countries is characterized by the contribution of 36.3 percent of all abstracts by United States (Fig. 1d). Around 5 percent of articles each are published from United Kingdom, Italy, Japan, Germany and Canada. 23 percent of all articles could not be matched to a country. It seems unlikely that those high publishing countries are specialized to research on cell death related topics, instead, as was also observed in previous studies, it rather reflects the development status of the respective country [12].

### Linguistic characteristics

Abstracts were divided into sentences, with a mean of 6.8 (sd ±2.86) and further into tokens, and token lemmas. The corpus has a length of 33,826,140 tokens and contains 188,052 lemmas, with a mean abstract length of about 180 tokens (sd ±77.4), the latter possibly due to the constraint on word length in abstract which is usually below 300 words for many journals.

#### Number of words per sentence differs across countries

Sentences comprise about 23words in average. A tendency to write longer sentences can be observed for Italy, whereas abstracts from China and Netherlands are made up of slightly shorter sentences in average (Fig. 2 & Table 1).

#### Lexical diversity is invariable across countries

Lexical diversity was assessed via type-token ratio calculated from 100 fold sampling of 100 abstracts for each country, thus avoiding that frequently publishing countries being represented by a higher number of experts on more diverse topics are seemingly characterized by higher lexical diversity due to a greater variety of technical terms needed.

**Fig 2.**
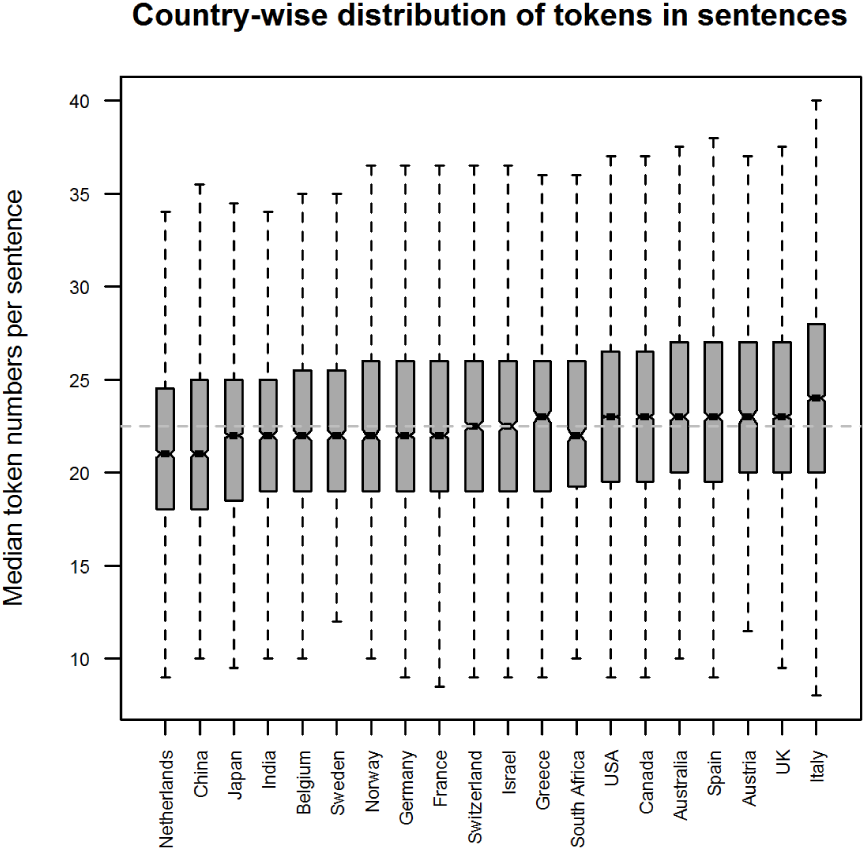
Tokens-per-sentence distribution across most-publishing countries varies only little. Abstracts from Netherlands and China tend to be composed of shorter sentences whereas abstracts from Italy are represented with longer sentences.

Interestingly, in spite of using English language as a secondary means of communicating research for not-native speakers, lexical diversity is invariantly maintained across countries (Table 1), and 17–18% lemma types are used in combination to form all tokens in the abstracts. In other words, of 100 tokens in an abstract, only 18 are unique, and the remaining 82 are just different forms of those unique words.

**Table 1.**
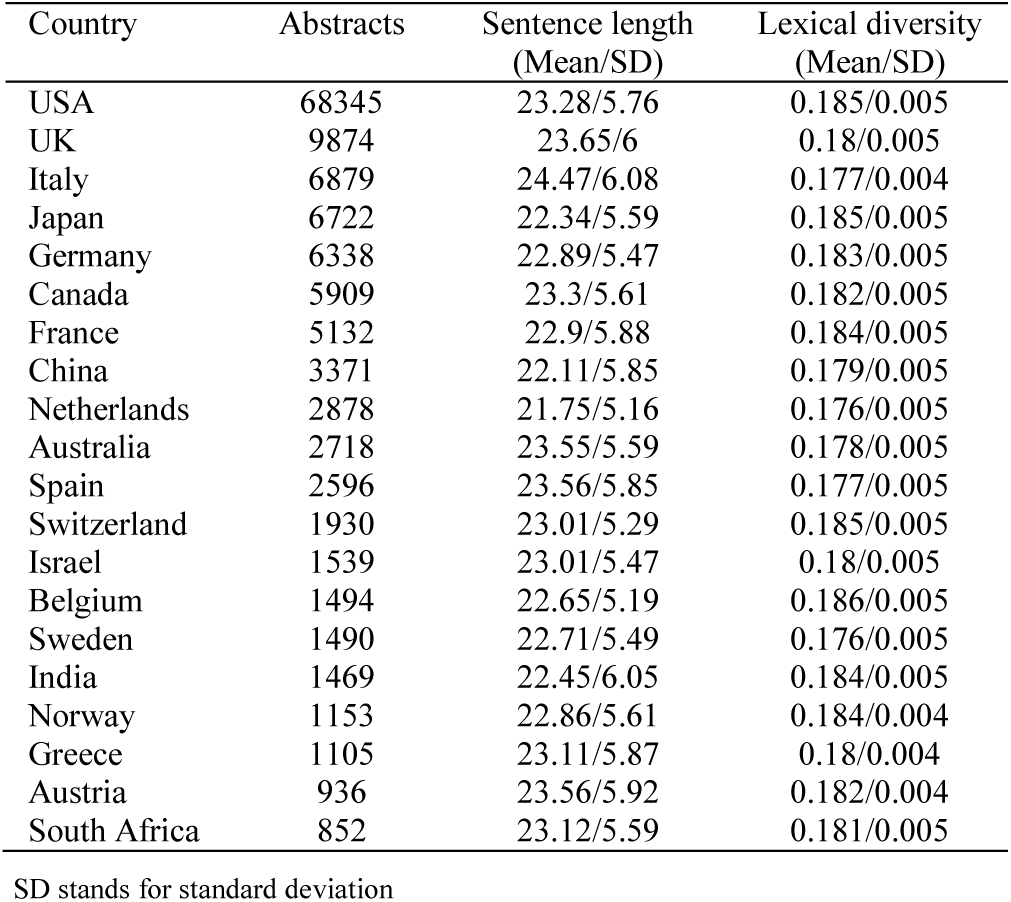
Country-wise sentence length and lexical diversity

| Country | Abstracts | Sentence length<br>(Mean/SD) | Lexical diversity<br>(Mean/SD) |
| --- | --- | --- | --- |
| USA | 68345 | 23.28/5.76 | 0.185/0.005 |
| UK | 9874 | 23.65/6 | 0.18/0.005 |
| Italy | 6879 | 24.47/6.08 | 0.177/0.004 |
| Japan | 6722 | 22.34/5.59 | 0.185/0.005 |
| Germany | 6338 | 22.89/5.47 | 0.183/0.005 |
| Canada | 5909 | 23.3/5.61 | 0.182/0.005 |
| France | 5132 | 22.9/5.88 | 0.184/0.005 |
| China | 3371 | 22.11/5.85 | 0.179/0.005 |
| Netherlands | 2878 | 21.75/5.16 | 0.176/0.005 |
| Australia | 2718 | 23.55/5.59 | 0.178/0.005 |
| Spain | 2596 | 23.56/5.85 | 0.177/0.005 |
| Switzerland | 1930 | 23.01/5.29 | 0.185/0.005 |
| Israel | 1539 | 23.01/5.47 | 0.18/0.005 |
| Belgium | 1494 | 22.65/5.19 | 0.186/0.005 |
| Sweden | 1490 | 22.71/5.49 | 0.176/0.005 |
| India | 1469 | 22.45/6.05 | 0.184/0.005 |
| Norway | 1153 | 22.86/5.61 | 0.184/0.004 |
| Greece | 1105 | 23.11/5.87 | 0.18/0.004 |
| Austria | 936 | 23.56/5.92 | 0.182/0.004 |
| South Africa | 852 | 23.12/5.59 | 0.181/0.005 |

SD stands for standard deviation

### Human migration across the globe might have influence on grammatical rules in writing

As an indicator of typical sentence structuring, POS-tag frequency tables containing 36 tags excluding punctuation were used to compare and cluster countries (Fig. 3)

**Fig 3.**
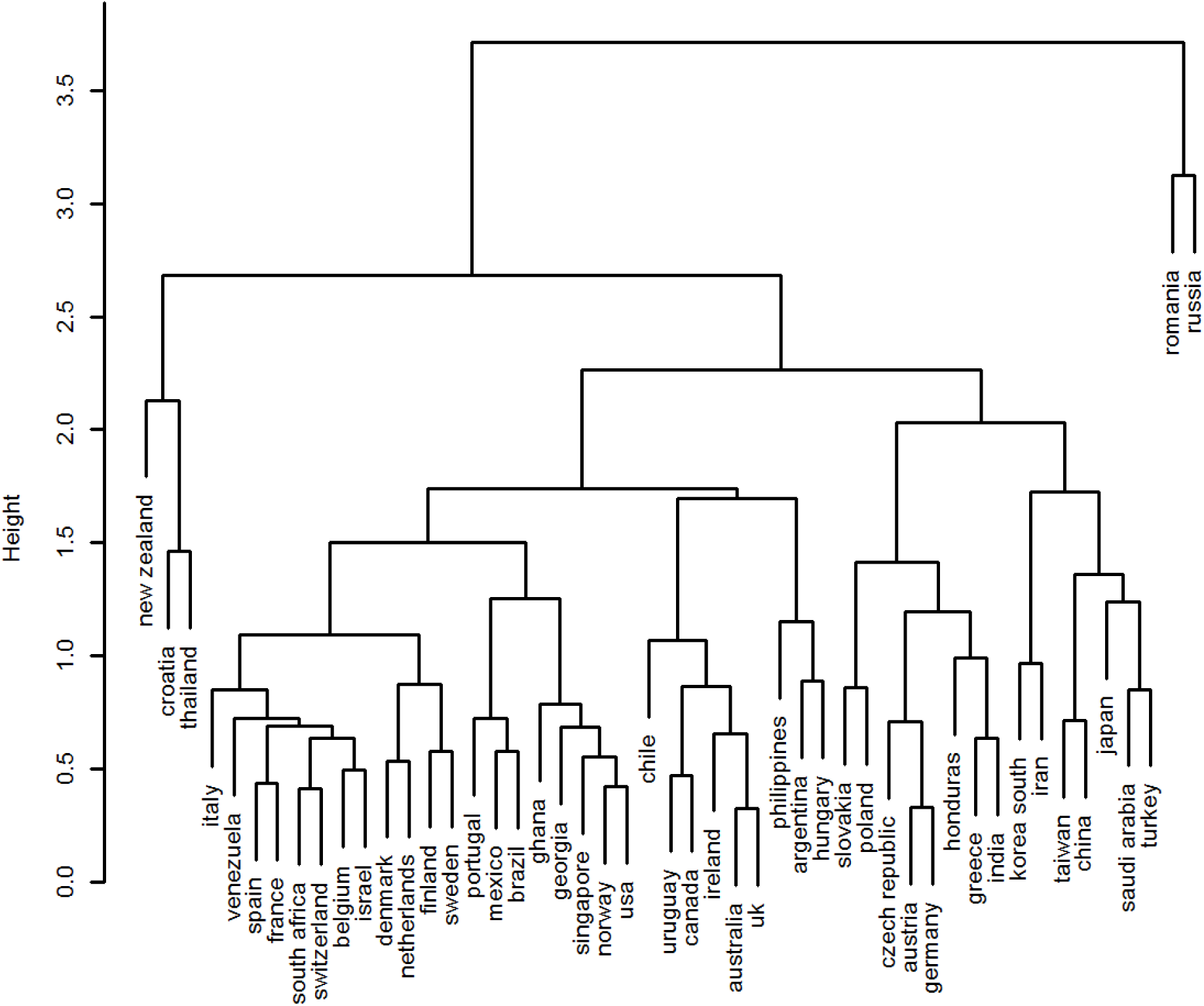
Linguistic style is similar for countries speaking same languages. Hierarchical clustering of countries based on Part-of-Speech-tag frequencies. Countries with language contact due to historical relations or spatial vicinity tend to cluster together. However, differences are subtle and result in low dendrogram height.

In several instances, the dendrogram suggests that the writing style in English is influenced by the mother tongue background of authors; for example France, Switzerland and Belgium are clustered together and share French as (one) official language. For researchers based in Canada, Ireland, Australia and the United Kingdom, it can be assumed that a considerable fraction is mother tongue speakers of English, and those countries, too, cluster together. These results are consistent with previous study [13]. Also, neighboring countries that share a border often cluster together, such as Slovakia, Poland, the Czech Republic, Austria and Germany, even though native languages may be different. Scientific writing in English has similar characteristics across those countries, suggesting that human migration and the associated transfer of native language background also plays a role in developing the writing style. This is supported by certain clusters that might hint at historic relations and social structures that have left their mark in people’s language, such as South Africa being clustered with Netherlands and Belgium.

In contrast to that, there seems to be no obvious relation between New Zealand, Croatia, and Thailand forming a separate cluster; and no clear reason why New Zealand should cluster so far from Australia while other spatially rather close countries often share a branch, such as is the case with Japan, China and South Korea. A possible reason may be that the differences in writing style are quite subtle, as may be concluded from the low height of the dendrogram, and thus noise might have considerable impact on the clustering.

In general, style differences seem subtle, as can be inferred from the low dendrogram height, however this may be contributed to by low impact research that might be published in less common journals that allow publications in regional languages as well. The limitation to English abstracts thus may favour the rather momentous publications by exceptionally well educated authors of countries with official and minority languages other than English.

## 4 Conclusions

The topic of cell death has been investigated thoroughly through the past five decades and contributions have been made by countries from all over the world. The publishing frequency however suggests that science and research still is a privilege to and widely dominated by developed countries. This is also reflected by the most publishing journals, many of which have a long standing history of publications addressing the topic of cell death.

Clustering of the most contributing countries with respect to the frequency of POS tags as an indication of writing style reveals only subtle differences that may be accounted for by native language background.

## 5 Acknowledgements

We acknowledge Dr. Vishal Acharya for critical reading of the manuscript.

## References

[1] J. Hirschberg and C. D. Manning, “Advances in natural language processing,” Science, vol. 349, pp. 261–6, Jul 17 2015.

[2] E. Brill, “A simple rule-based part of speech tagger,” presented at the Proceedings of the third conference on Applied natural language processing, Trento, Italy, 1992.

[3] U. Hahn and J. Wermter, “Tagging Medical Documents with High Accuracy,” in PRICAI 2004: Trends in Artificial Intelligence. vol. 3157, C. Zhang, H. W. Guesgen, and W.-K. Yeap, Eds., ed: Springer Berlin Heidelberg, 2004, pp. 852–861.

[4] G. Salton and M. J. McGill, Introduction to Modern Information Retrieval: McGraw-Hill, Inc., 1986.

[5] M. Bates, “Models of natural language understanding,” Proc Natl Acad Sci U S A, vol. 92, pp. 9977–82, Oct 24 1995.

[6] E. W. Sayers, T. Barrett, D. A. Benson, E. Bolton, S. H. Bryant, K. Canese, et al., “Database resources of the National Center for Biotechnology Information,” Nucleic Acids Res, vol. 40, pp. D13–25, Jan 2012.

[7] H. Schmid, “Probabilistic Part-of-Speech Tagging Using Decision Trees,” Proceedings of International Conference on New Methods in Language Processing, Manchester, UK, 1994.

[8] I. Feinerer, K. Hornik, and D. Meyer, “Text Mining Infrastructure in R,” 2008, vol. 25, p. 54, 2008–03–31 2008.

[9] J. E. Prier and R. S. Brodey, “Canine Neoplasia. A Prototype for Human Cancer Study,” Bull World Health Organ, vol. 29, pp. 331–44, 1963.

[10] D. J. Klionsky, H. Abeliovich, P. Agostinis, D. K. Agrawal, G. Aliev, D. S. Askew, et al., “Guidelines for the use and interpretation of assays for monitoring autophagy in higher eukaryotes,” Autophagy, vol. 4, pp. 151–75, Feb 2008.

[11] L. Lelievre, D. Garcia-Hermoso, H. Abdoul, M. Hivelin, T. Chouaki, D. Toubas, et al., “Posttraumatic mucormycosis: a nationwide study in France and review of the literature,” Medicine (Baltimore), vol. 93, pp. 395–404, Nov 2014.

[12] F. Salager-Meyer, “Scientific publishing in developing countries: Challenges for the future,” Journal of English for Academic Purposes, vol. 7, pp. 121–132, 4// 2008.

[13] R. Netzel, C. Perez-Iratxeta, P. Bork, and M. A. Andrade, “The way we write,” EMBO Rep, vol. 4, pp. 446–51, May 2003.

